# Celecoxib exhibits therapeutic potential in experimental model of hyperlipidaemia

**DOI:** 10.1101/2021.02.15.431245

**Authors:** Martins Ekor, Phyllis E. Owusu Agyei, Ernest Obese, Robert P. Biney, Isaac T. Henneh, Meshack Antwi-Adjei, Ewura S. Yahaya, Gordon Amoakohene, Patrick K. Akakpo

**Author notes:** Corresponding Author: and (ME).

## Abstract

Hyperlipidaemia is a major risk factor for cardiovascular diseases, the leading cause of death globally. Celecoxib attenuated hypercholesterolemia associated with CCl_4_-induced hepatic injury in rats without improving liver function in our previous study. This present study investigated the lipid lowering potential of celecoxib in normal rats fed with coconut oil subjected to five deep-frying episodes. Male Sprague Dawley rats were randomly assigned to groups (n=6 rats/group) which received physiological saline (10 mL/kg), unheated coconut oil (UO, 10 mL/kg) or heated coconut oil (HO, 10 ml/kg) for 60 days. Groups that received HO were subsequently treated with either physiological saline, atorvastatin (25 mg/kg), celecoxib (5 mg/kg) or celecoxib (10 mg/kg) in the last fifteen days of the experiment. Rats were sacrificed 24 hours after last treatment and blood and tissue samples collected for analysis. HO consumption produced significant hyperlipidaemia and elevation in marker enzymes of hepatic function. Celecoxib ameliorated the hyperlipidaemia as shown by the significantly (P<0.05) lower total cholesterol, triglycerides, low and very low density lipoprotein in the celecoxib-treated rats when compared with HO-fed rats that received saline. Celecoxib also reduced (P<0.05) alanine aminotransferase, aspartate aminotransferase, alkaline phosphatase and liver weight of hyperlipidaemic rats. Similarly, hepatocellular damage and inflammation of the aorta associated with the hyperlipidaemia was significantly reversed by celecoxib. However, serum TNF-α and IL-6 did not change significantly between the various groups. Taken together, data from this study suggest that celecoxib may exert therapeutic benefit in hyperlipidaemia and its attendant consequences.

## Introduction

Hyperlipidaemia, also recognised as dyslipidaemia, describes the manifestation of different disorders of lipoprotein metabolism [1]. Patients with hyperlipidaemia are mostly asymptomatic but have an increased risk for cardiovascular diseases (CVDs). CVDs are recognized as one of the leading causes of mortality and a major cause of morbidity worldwide [2–6]. Atherosclerosis, a vascular disease affecting blood circulation in the coronary, central, and peripheral arteries, is the major form of CVD and it is characterized by chronic inflammatory build up, driven largely by lipid accumulation within the walls of the artery. Unlike acute inflammation, atherosclerosis is hallmarked by a state of unresolved low-grade chronic inflammation. Importantly, low-grade inflammation is also a feature of several diseases known to increase the risk of CVD [3]. Besides hypertension, chronic dyslipidaemia is a major cause of atherosclerosis [7]. Although elevated low density lipoprotein cholesterol (LDL-C) is thought to be the best indicator of atherosclerosis risk, dyslipidaemia can also describe elevated total cholesterol (TC) or triglycerides (TG), or low levels of high density lipoprotein cholesterol (HDL-C) [8].

Hyperlipidaemia may affect the severity of tissue damage in other pathological conditions, notably in liver injury [9]. Primary associated clinical findings of fatty liver are hyperlipidaemia, hyperglycaemia, hypertension, and hyperuricemia [10]. Although, some success has been achieved with the use of statins in the management of hyperlipidaemia, with reports of improved quality of life and decreased mortality and morbidity in many patients with CVDs [11], the use of statins has been associated with side effects such as myopathy, headache, bowel upset, nausea, sleep disturbance, increased creatinine phosphokinase and serum transaminase hence requiring routine monitoring of these parameters [12]. Fibrates, bile acid sequestrants and nicotinic acid which constitute other modalities of treatment have some side effects as well [12]. This notwithstanding, their control of lipid levels is far from satisfactory. This calls for increased search for newer drugs with hypolipidaemic properties or repurposing of existing drugs for use in hyperlipidaemic conditions.

The non-steroidal anti-inflammatory drugs (NSAIDs) are used primarily for the management of inflammatory conditions such as arthritis and are known to exert their effect via inhibition of cyclooxygenase (COX-1 and COX-2) activity [13]. At higher concentrations, NSAIDs are also known to reduce production of superoxide radicals, induce apoptosis, impede the expression of adhesion molecules, decrease nitric oxide synthase, decrease proinflammatory cytokines (*e.g.,* TNF-α, interleukin-1), modify lymphocyte activity and alter cellular membrane functions [14]. All these markers are known to be up-regulated in inflammatory conditions and other disorders which have inflammatory subsidiaries. NSAIDs such as ibuprofen have been shown to lower plasma cholesterol levels and reduce the progression of atherosclerosis in humans and laboratory animals [15–17]. Other studies have shown that indomethacin lowers the cholesterol content in liver and atherosclerotic blood vessels [18, 19]. Most of these effects of NSAIDs have been observed in *in vitro* cell cultures, or with atherogenic diets in rabbits [13].

The Coxibs, designed to selectively block COX-2, appeared a promising solution in the effort to avoid the gastrointestinal and other adverse effects that were noted with traditional NSAIDs [20, 21]. Celecoxib was the first specific COX-2 inhibitor to be approved for the treatment of rheumatic diseases. Observations from several clinical studies have led to concerns being raised about the cardiovascular safety of the COX-2 inhibitors. While the evidence regarding the cardiovascular risk associated with these drugs was not encouraging, a number of studies demonstrated that celecoxib is safer than other coxibs. Several preclinical and clinical studies have shown that celecoxib is capable of exerting a beneficial impact on cardiovascular health [22–25].

There has been a general assumption that COX-2 inhibitors may be beneficial in atherosclerosis, liver disease and hypercholesterolaemia since pathogenesis of these diseases is closely linked with prostaglandins [15] and since upregulation of COX-2 expression has also been demonstrated in hyperlipidaemia [26]. In our previous study, celecoxib was observed to significantly attenuate hypercholesterolemia and lipid peroxidation associated with liver injury during carbon-tetrachloride–associated hepatotoxicity in rats [27]. This important observation needs further evaluation in experimental hyperlipidaemia models devoid of hepatotoxin to ascertain the possible therapeutic potential of celecoxib in hyperlipidaemia. Considering the huge cost, time, safety and legal challenges associated with discovery of newer drugs, repurposing of already existing drugs in clinical use for newer indications provides rapid alternative to ensure improved access to medicines with relatively minimal resources. It is on this basis that we evaluate the FDA-approved selective COX-2 inhibitor, celecoxib, as a potential addition to the already existing pharmacotherapy for hyperlipidaemia in the current study.

## Materials and methods

### Animals

Male Sprague-Dawley rats (170-250 g) were obtained from the Noguchi Memorial Institute for Medical Research, Ghana. The animals were housed in stainless cages (34 × 47 × 18) in groups of five at the animal house facility of School of Biological Sciences, University of Cape Coast. Animals were fed with normal commercial diet bought from Flour Mills of Ghana Limited, Tema, Ghana and water was provided *ad libitum*. They were kept under normal laboratory conditions with regards to room temperature and humidity. All the techniques and protocols used in the study were done in accordance with established public health guidelines in “Guide for Care and Use of Laboratory Animals” [28] (Garber *et al*. 2011).

### Drugs and chemicals

Celecoxib (Celebrex™) and Atorvastatin (Lipitor^®^) were purchased from Pfizer Pharmaceutical LLC, Vega Baja, Puerto Rico Virgin^®^ coconut oil was purchased from the Kotokuraba market at Cape Coast, Ghana.

### Experimental design and treatment

Thirty-six rats (weighing between 170–250 g) were divided into six groups of 6 rats per group and fed with normal commercial diet during the 7-day acclimatization period and throughout the 60-day experimental period. Animals were treated as follows as shown in the Table 1. The doses of atorvastatin (ATO) and celecoxib (CXB) were selected according to previous studies [27]. Animals were fasted overnight after all the treatments and fasting blood glucose levels were measured 24 h after the final treatment. The animals were humanely sacrificed by cervical dislocation and blood and organs were harvested for other investigations. Blood samples were collected via cardiac puncture into EDTA and gel separator tubes for haematological and biochemical analyses, respectively.

**Table 1:**
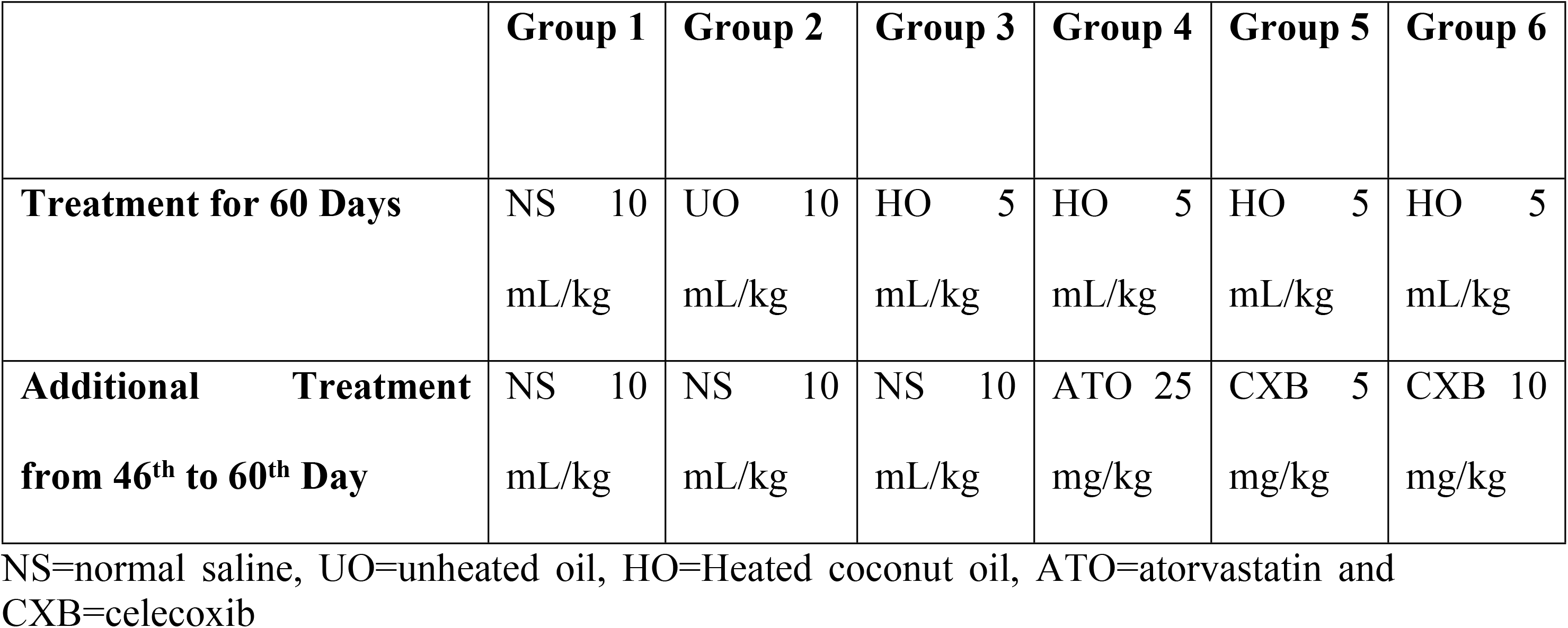
Treatment schedule.

### Relative weight of organs

Specific organs, including the liver, heart, kidney, lungs and spleen were harvested and weighed. Relative organ weights (mg/kg body weight) were estimated and values analyzed.

### Haematological analysis

Blood samples were analyzed by haem automated analyzer (CELL-DYN 1700, Abbot Diagnostics Division, Abbot Laboratories, Abbot Park, Illinois, USA) for total blood count and specific differentials.

### Biochemical analysis

Blood samples were allowed to clot for 30 min at room temperature and centrifuged at 1000 rpm for 10 min. Serum obtained was stored at −20°C until biochemical analysis was carried out. Serum indices were analyzed by an automated analyzer (ATAC 8000 Random Access Chemistry System, Elan Diagnostics, Smithfied, RI, USA) and estimations for aspartate aminotransferase (AST), alanine aminotransferase (ALT), alkaline phosphatase (ALP), Creatinine, blood urea nitrogen (BUN), fasting blood sugar (FBS), total cholesterol (TC), triglycerides (TG), high density lipoprotein (HDL) cholesterol direct, low-density lipoprotein (LDL) cholesterol and very low-density lipoprotein (VLDL) were recorded

### Serum Cytokine (IL-6 and TNF-α) levels

The blood samples were centrifuged at 1000 rpm for 10 min. Sera formed were aliquoted into eppendorf tubes and stored at −20°C before analysis. Serum levels of IL-6 and TNF-α were estimated in duplicates with specific rat ELISA kit (Boster Biological Technology 3942 Valley Ave Pleasanton, CA 94566, USA) assay in accordance with the recommendations of the manufacturer. The absorbance of the samples was read at 450 nm using a micro-plate spectrometer (Spectramax 190 Micro-plate Spectrometer, 90-250V 50-60Hz, Molecular Devices, CA, USA).

### Histopathological studies

Portions of the tissues from liver, kidney, heart, lungs, spleen and aorta were used for histopathological examination. Tissues were fixed in 10% neutral buffered formalin (pH 7.2) and dehydrated through a series of ethanol solutions, embedded in paraffin and routinely processed for histological analysis. A section (2 μm thickness) was cut and stained with haematoxylin-eosin for examination. The stained tissues were observed through an Olympus BX-51 microscope (Olympus Corporation, Tokyo, Japan) and photographed by INFINITY 4 USB Scientific Camera (Lumenera Corporation, Otawa, Canada).

### Data analysis

Data has been presented as mean of six rats ± standard error of mean (SEM). The presence of significant differences between means of groups was determined by one-way analysis of variance (ANOVA) using GraphPad Prism for Windows version 7 (GraphPad Software, San Diego, CA, USA). Significant difference between groups was determined using the Newman-Keuls’ Multiple Comparison Test with P<0.05 considered statically significant.

## Results

### Changes in relative organ weights

The results presented in Fig 1 describe the effect of celecoxib (CXB 5 and 10 mg/kg) and atorvastatin (ATO 25 mg/kg) on hyperlipidaemia induced by heated coconut oil. Naïve control group received only normal saline (10 mL/kg) throughout the experiment, the negative control group received heated oil and normal saline (10 mL/kg) whereas another group received unheated oil and normal saline (10 mL/kg). The relative liver weight (liver-to-body weight ratio) of the group that received only heated oil was significantly (P < 0.05) higher compared to the naïve control and unheated oil group. This was, however, significantly (P<0.05) decreased by treatment with celecoxib and atorvastatin. The relative weights of other organs such as heart, kidney, lungs and spleen were not significantly affected as shown in Fig 1.

**Figure 1.**
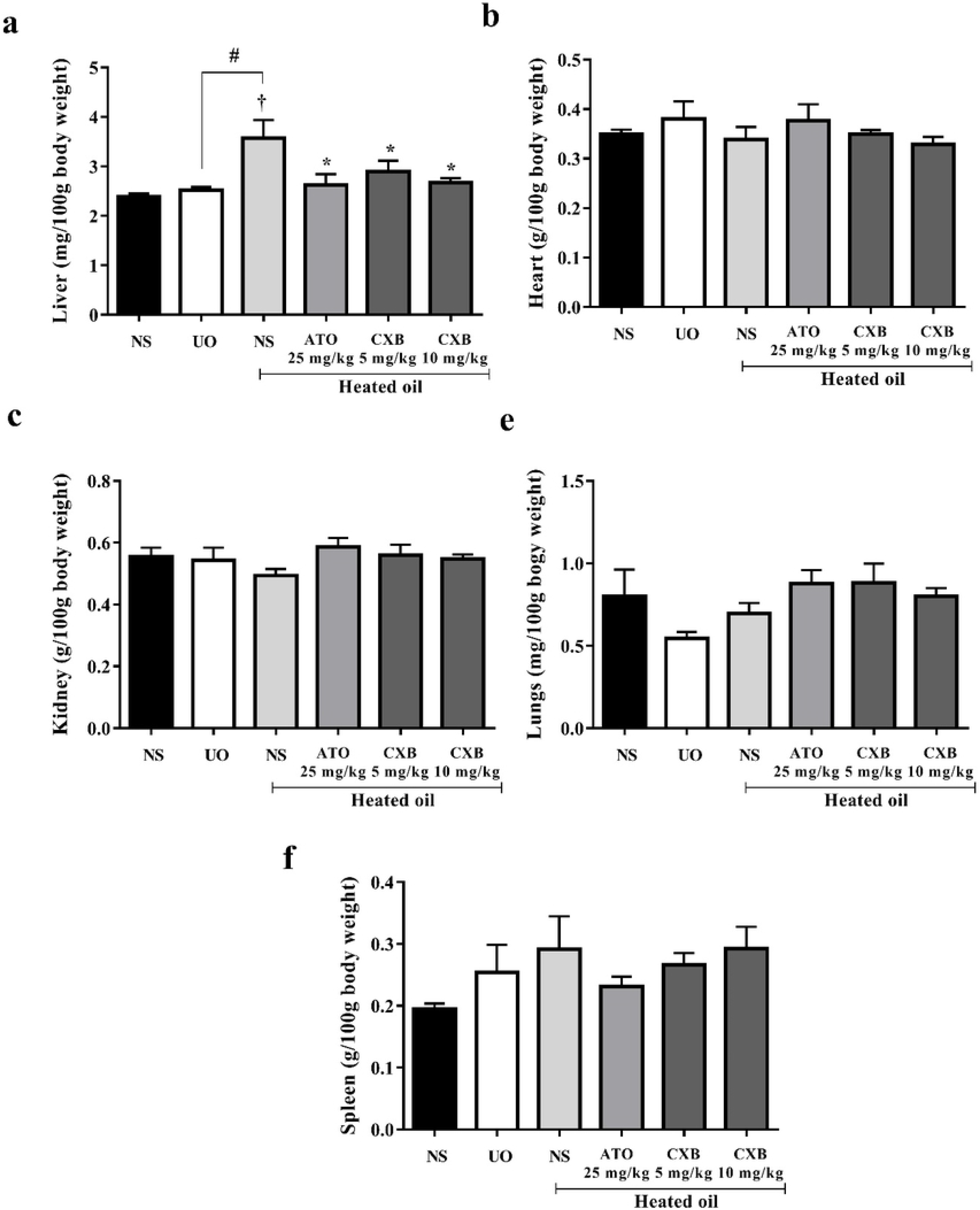
Effect of celecoxib (CXB 5 and 10 mg/kg), atorvastatin (ATO 25 mg/kg) on the weight of (a) liver, (b) heart (c) kidney (d) lungs and (e) spleen in overheated-oil induced hyperlipidemia in Sprague-Dawley rats. Values are expressed as mean ± SEM (n=6). The symbols * represents significant differences between treatment groups and heated oil only group (*P<0.05)*; *#* represents significant differences (*P<0.05)* between heated and unheated oil whereas † represents significant differences (*P<0.05)* between heated oil and naïve control group (all were compared using one-way ANOVA followed by Neuman Keals’ *post hoc* test).

### Changes in haematological parameters

The heated oil and the various drug treatments did not significantly alter haematological parameters such as the red blood cell count, haemoglobin, haematocrit, mean cell volume, mean cell haemoglobin concentration, platelet count, white blood cell count and its differentials as presented in Figs 2 and 3. However, there was a remarkable (P<0.05) increase in the platelet count (thrombocytosis) compared to the naïve control. This increase was also observed in the unheated oil treated group, though not significant (P>0.05) (Figs 2 and 3).

**Figure 2.**
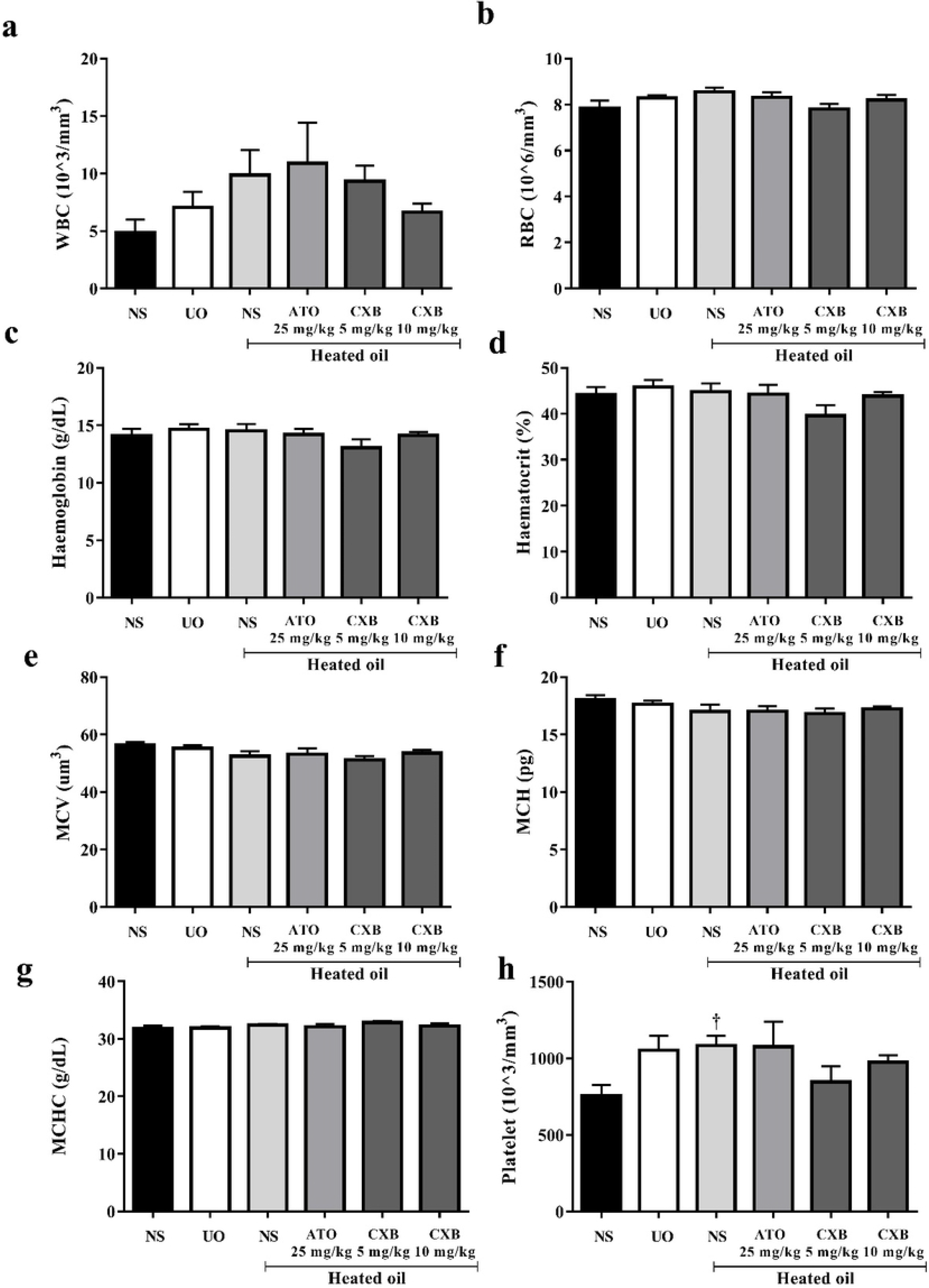
Effect of celecoxib (CXB 5 and 10 mg/kg), atorvastatin (ATO 25 mg/kg) on haematological parameters such as (a) white blood cells (b) red blood cells (RBCs) (c) haemoglobin (d) haematocrit (e) mean cell volume (MCV) (f) mean cell haemoglobin (g) mean cell haemoglobin concentration (MCHC) and (h) platelet in overheated-oil induced hyperlipidemia in Sprague-Dawley rats. Values are expressed as mean ± SEM (n=6). The † represents significant differences (*P<0.05)* between heated oil and naïve control group (compared using one-way ANOVA followed by Neuman Keals’ *post hoc* test).

**Figure 3.**
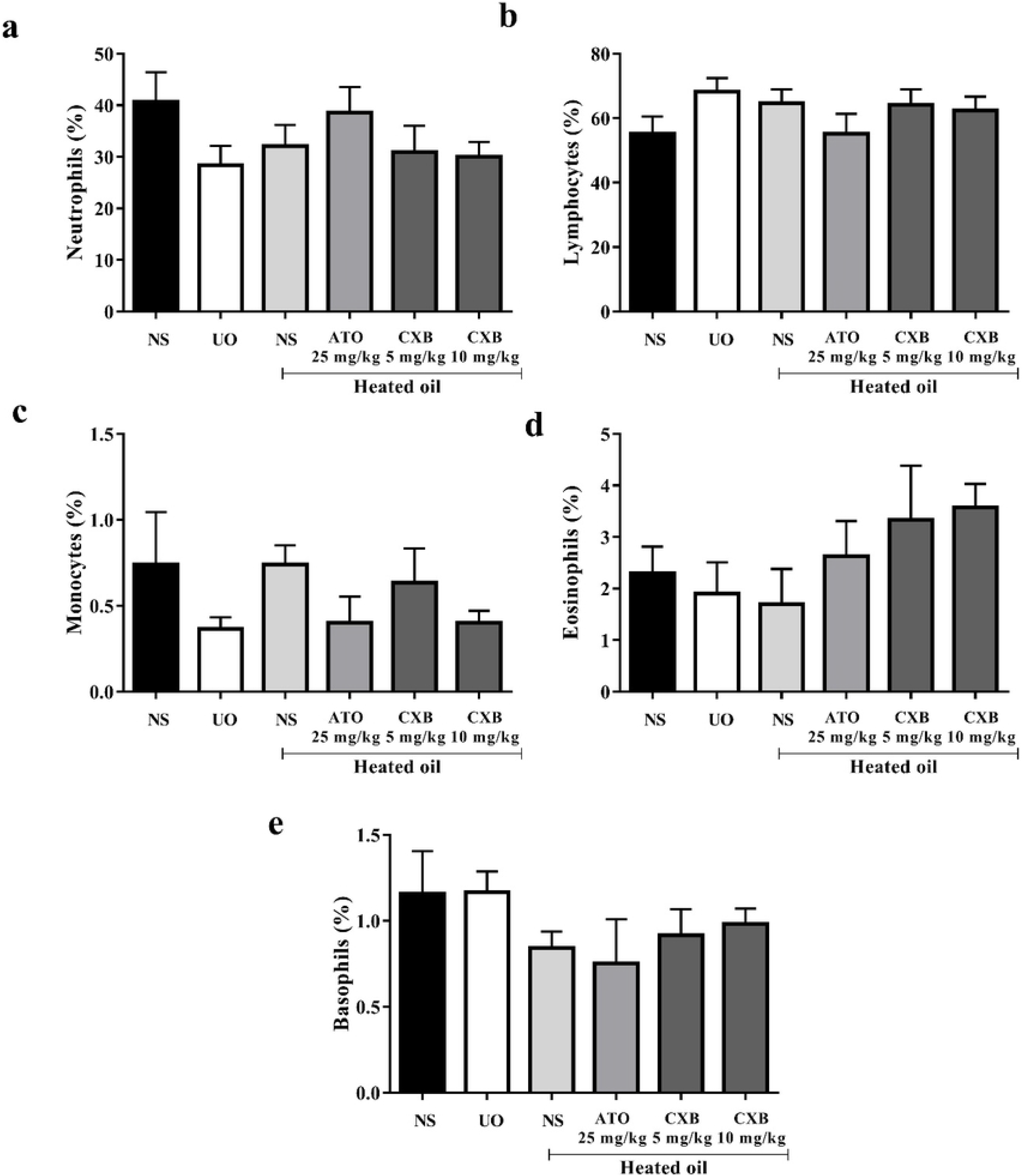
Effect of celecoxib (CXB 5 and 10 mg/kg), atorvastatin (ATO 25 mg/kg) on differential white blood cell parameters such as (a) neutrophils (b) lymphocytes (c) monocytes (d) eosinophils (e) basophils in overheated-oil induced hyperlipidemia in Sprague-Dawley rats. Values are expressed as mean ± SEM (n=6). There were no significant differences between treatment groups and the various controls (all were compared using one-way ANOVA followed by Neuman Keals’ *post hoc* test).

### Changes in serum biochemical parameters

Results presented in Fig 4 show alanine aminotransferase (ALT) and alkaline phosphatase (ALP) activities of rats fed with heated coconut oil were significantly (P<0.05) higher than those of the naïve control. However, treatment with celecoxib (5 and 10 mg/kg) significantly (P<0.01) reversed these elevations in the liver enzymes. Activities of aspartate aminotransferase (AST) enzyme as well as fasting blood glucose were however, not significant different among the various treatment groups.

**Figure 4.**
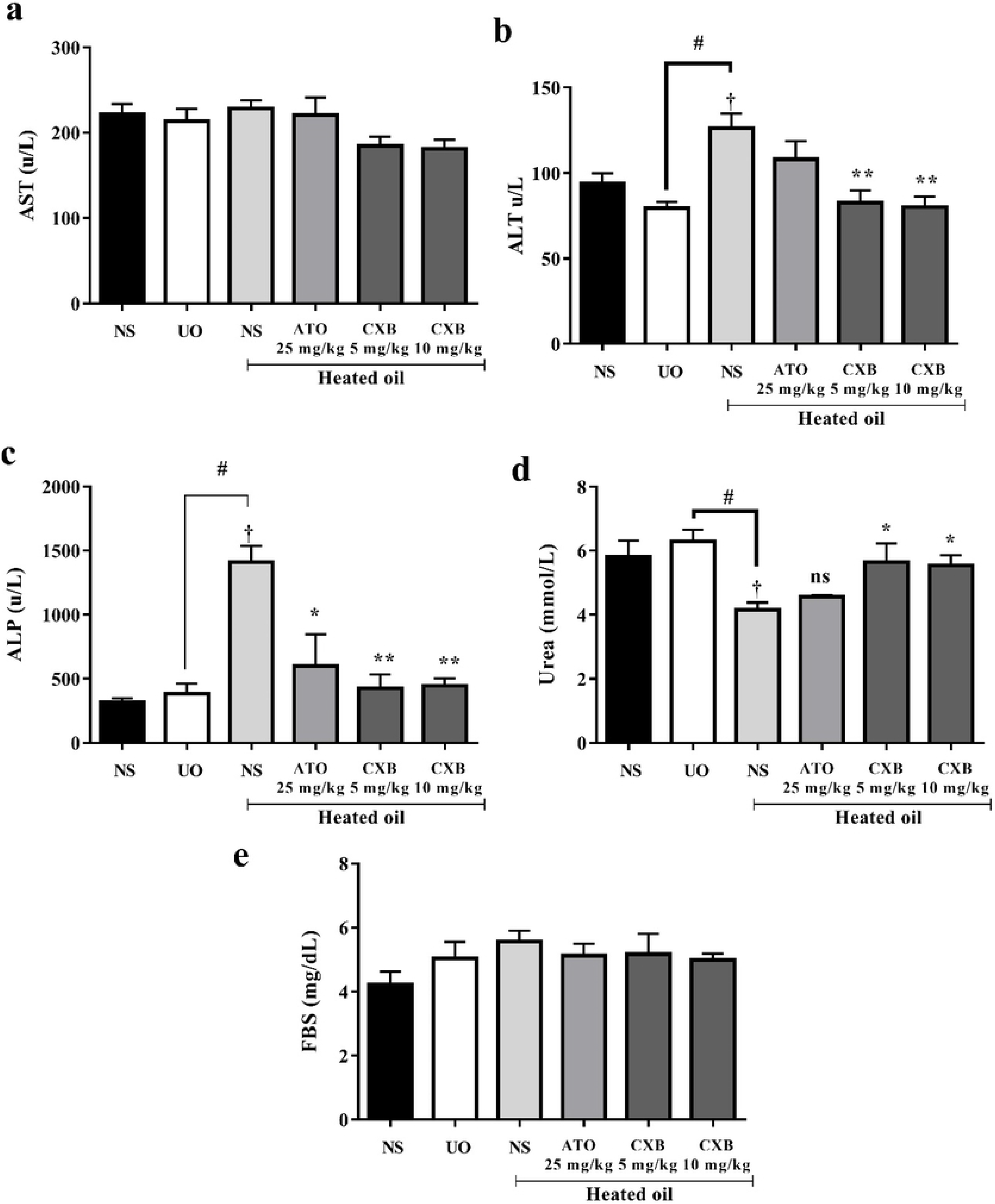
Effect of celecoxib (CXB 5 and 10 mg/kg), atorvastatin (ATO 25 mg/kg) on serum lipid parameters such as (a) AST (b) ALT (c) ALP and (d) urea; and (e) fasting blood sugar in overheated-oil induced hyperlipidemia in Sprague-Dawley rats. Values are expressed as mean ± SEM (n=6). The symbols * and ** represents significant differences (*P<0.05 and P<0.01* respectively) between treatment groups and heated oil only group; *#* represents significant differences (*P<0.05)* between heated and unheated oil whereas † represents significant differences (*P<0.05)* between heated oil and naïve control group (all were compared using one-way ANOVA followed by Neuman Keals’ *post hoc* test).

With respect to urea however, treatment of rats with heated oil significantly reduced its levels compared to the naïve control as those treated with unheated oil. The decrease was however, significantly (P<0.05) reversed to normal by both doses of celecoxib but not atorvastatin (Fig 4).

### Changes in lipid profile

Treatment of rats with heated oil only significantly (P<0.05) elevated the levels of cholesterol, triglycerides, LDL as well as VLDL. All there parameters were significantly reversed by treatment with atorvastatin and celecoxib. The levels of high density lipoproteins (HDL) was not significantly affected compared to the controls despite the fact that the levels decreased in rats treated with unheated oil only (Fig 5).

**Figure 5.**
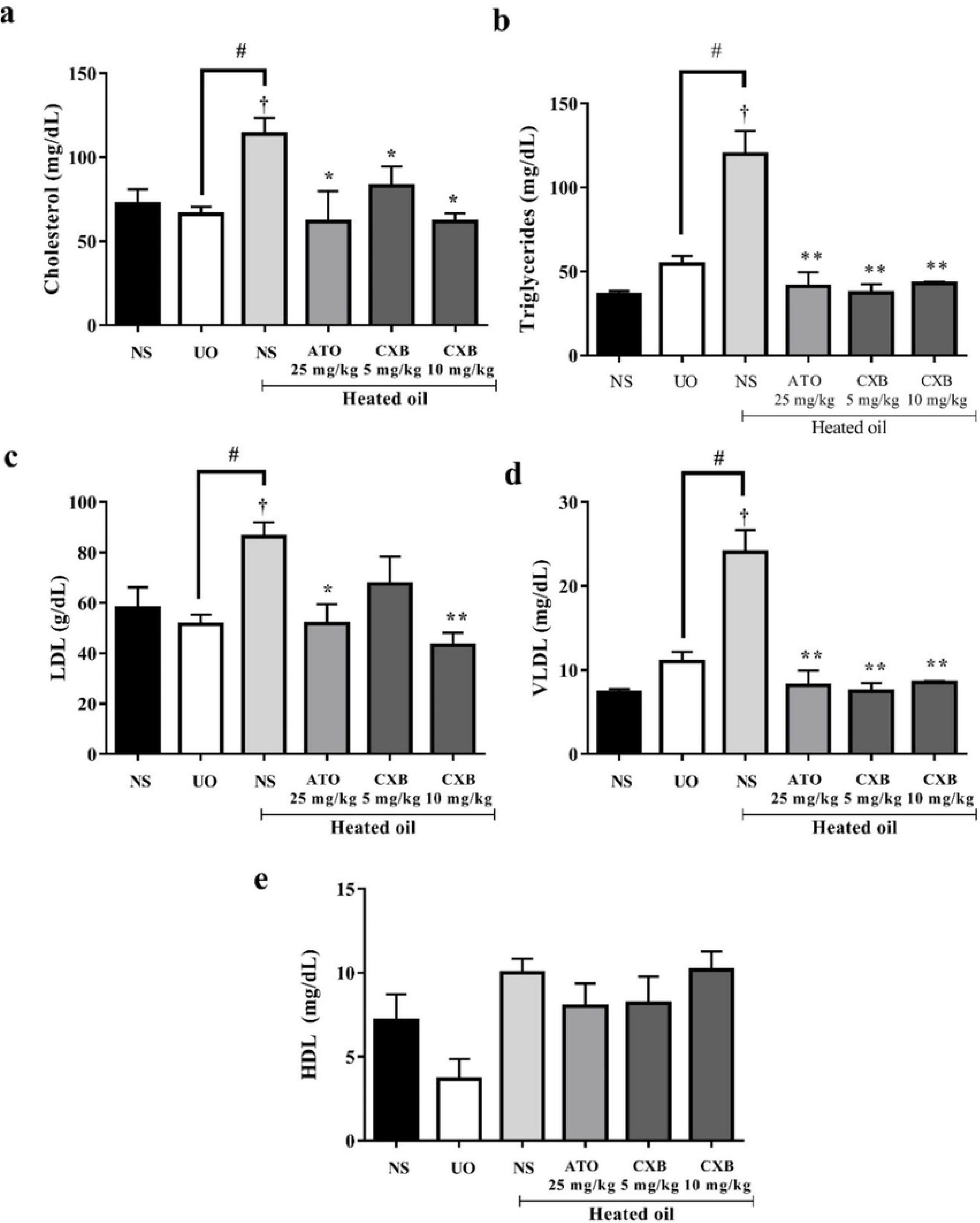
Effect of celecoxib (CXB 5 and 10 mg/kg), atorvastatin (ATO 25 mg/kg) on serum lipid parameters such as (a) cholesterol (b) triglycrides (c) low density lipoprotein (d) very low density lipoprotein and (e) high density lipoprotein cholesterol in overheated-oil induced hyperlipidemia in Sprague-Dawley rats. Values are expressed as mean ± SEM (n=6). The symbols * and ** represents significant differences (*P<0.05 and P<0.01* respectively) between treatment groups and heated oil only group; *#* represents significant differences (*P<0.05)* between heated and unheated oil whereas † represents significant differences (*P<0.05)* between heated oil and naïve control group (all were compared using one-way ANOVA followed by Neuman Keals’ *post hoc* test).

### Changes in cytokine levels

Results presented in Figure show that treatment of rats with heated coconut oil did not induce significant changes in the levels both TNF-α as well as interleukin 1β compared with naïve control group. Additionally, the celecoxib as well as atorvastatin did not alter the levels of the cytokines significantly (Fig 6).

**Figure 6.**
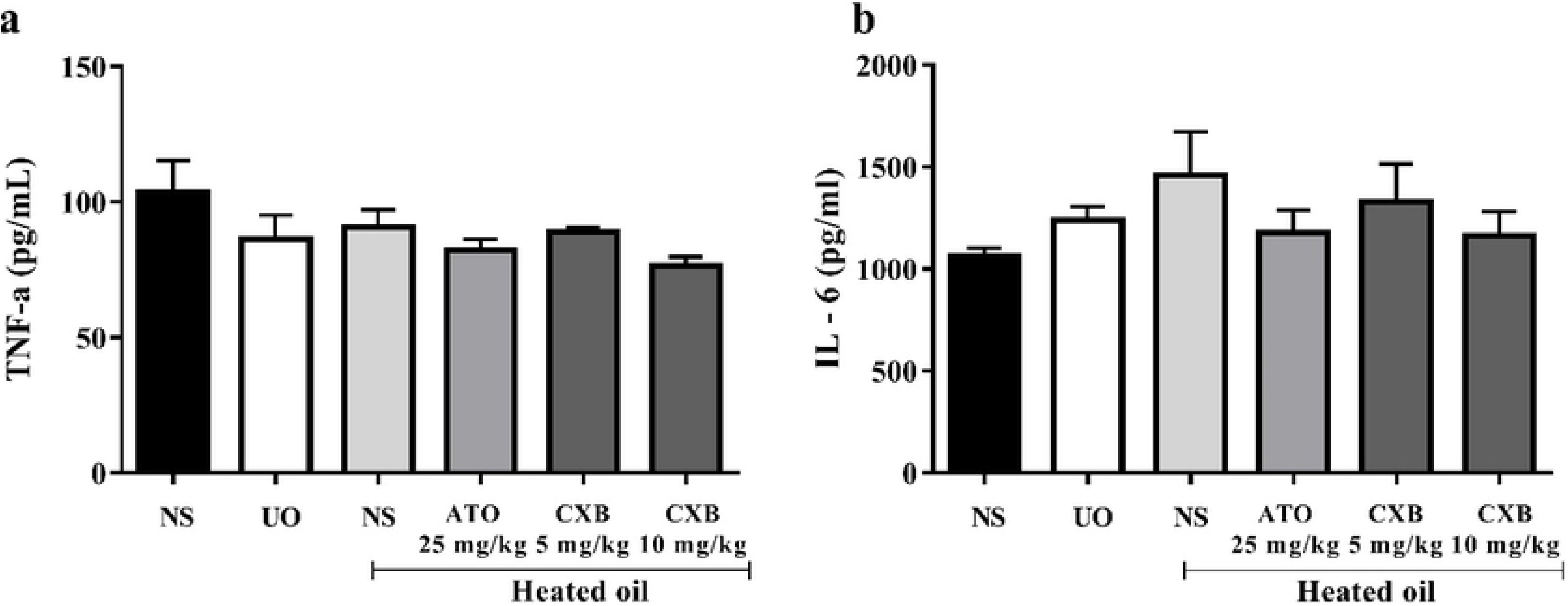
Effect of celecoxib (CXB 5 and 10 mg/kg) and atorvastatin (ATO 25 mg/kg) on the levels of serum cytokines such as (a) TNF-α and (b) IL-6 in overheated-oil induced hyperlipidemia in Sprague-Dawley rats. Values are expressed as mean ± SEM (n=6). There were no significant differences between heated oil and other treatment groups (all were compared using one-way ANOVA followed by Neuman Keals’ *post hoc* test).

### Histopathological changes in the liver and aorta

Photomicrographs shown in Fig 7 A contain a section of the liver of naïve control rats. The architecture of the liver is normal with congestion of central veins. The hepatocytes appear histologically normal without any evidence of degeneration, fatty change or necrosis. This is comparable to Fig 7 B which is section of a representative liver sample from rats that received unheated oil. The hepatocytes appear histologically normal without evidence of degeneration, fatty change or necrosis even though in some areas there are mild periportal chronic inflammatory infiltrates. Figure 7C shows a section of representative liver photomicrograph from the negative control group. Treatment with heated oil only resulted in a poor dehydration of the tissue. The architecture of the liver was distorted in some areas and preserved elsewhere. There was intense congestion of central veins. Also, there was a large area of confluent necrosis with bridging, demarcated by only surrounding residual fibrocollagenous meshwork and surrounding central veins. There was also a proliferation of bile ductules around this area of infarction. The periportal vessels and the area surrounding the central vein shows intense chronic inflammatory changes. This is a case of massive necrosis with ductular reaction and hepatitis as shown in Fig 7 C. Treatment with atorvastatin (ATO 25 mg/kg) restored the hepatocytes and histoarchitecture of the liver to normal as shown in Figure 7D. Although sections of the liver tissue showed poor dehydration in this group, the architecture of the liver was normal despite the congestion of the central veins. Overall, the hepatocytes appeared histologically normal without evidence of degeneration, fatty change or necrosis. With rats treated with low dose celecoxib (CXB 5 mg/kg), sections of liver showed poor dehydration of tissue. The architecture of the liver was normal with congestion of central veins. The hepatocytes appeared histologically normal without evidence of degeneration, fatty change or necrosis. In some areas, there were mild periportal chronic inflammatory infiltrates. However, the tissue could be said to be histologically normal and this possibly show evidence of recovery following sub-chronic administration of heated oil as shown in Fig 7 E. The high dose celecoxib (CXB 10 mg/kg), however, showed mild chronic hepatitis as shown in Fig 7 F. Sections of the liver showed poor dehydration of tissue. The architecture of the liver was normal with congestion of central veins and the hepatocytes appeared histologically normal without evidence of degeneration, fatty change or necrosis. In some areas, there were moderate periportal chronic inflammatory infiltrates dominated by groups of lymphocytes that formed aggregates in those areas. There were no associated bridging or evidences of fibrosis.

**Figure 7:**
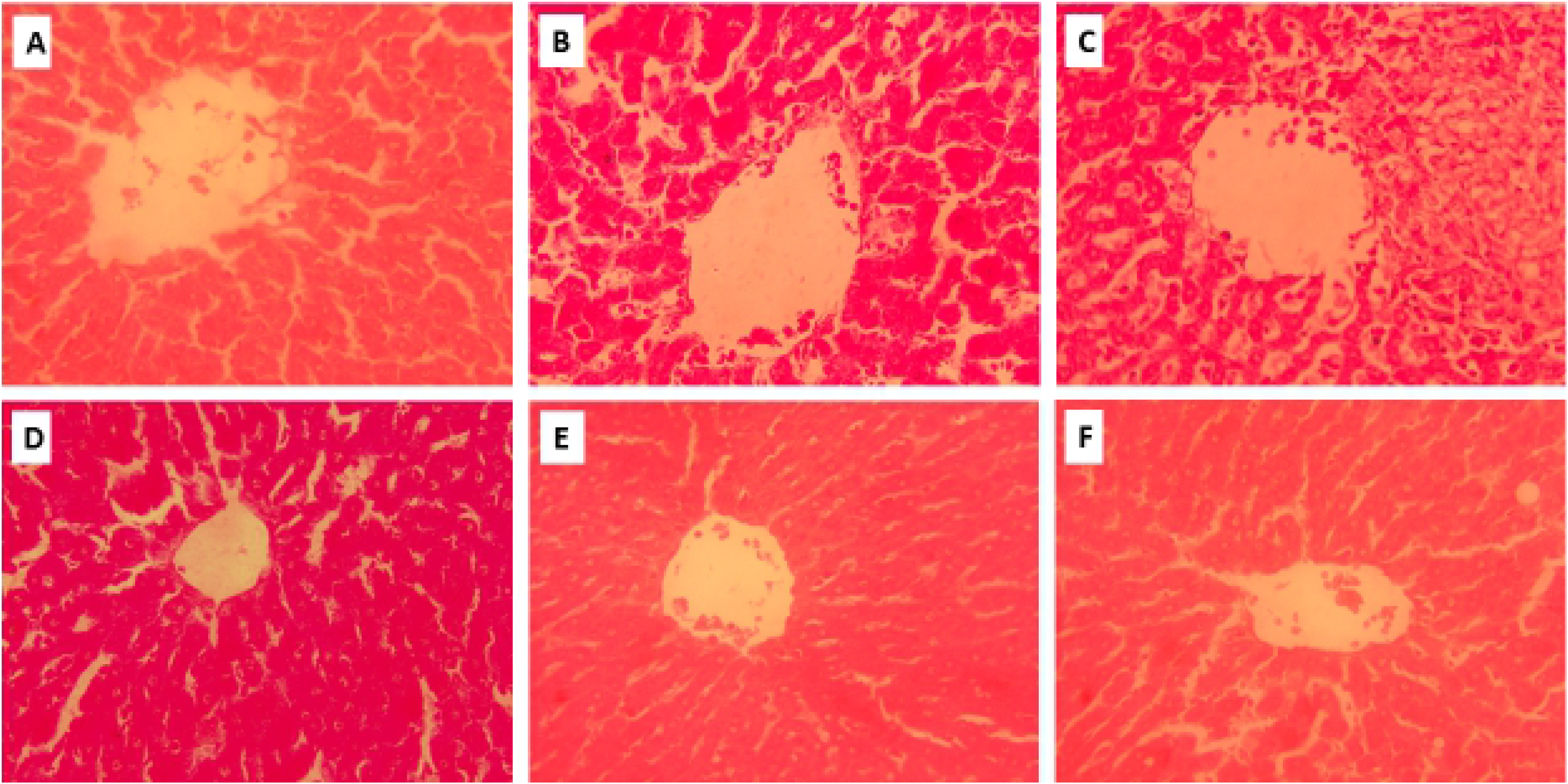
Photomicrograph of liver section of representative rat in (A) naïve control group (B) unheated oil treated group (C) heated oil only treated group as negative control, (D) heated oil in addition to atorvastatin 25 mg/kg (E) heated oil in addition to celecoxib 5 mg/kg and (F) heated oil in addition to celecoxib 10 mg/kg (H & E, ×40).

Photomicrographs presented in Fig 8 show the cross section of aorta of various representative samples from various treatment groups. Naïve control rats that received only normal saline without any drug treatment had a histologically normal aorta and perivascular tissues as shown in Fig 8 A. Sections show cross sections of the aorta with surrounding fibrofatty tissue. The lumen of the aorta has a collection of red blood cells with the wall of the aorta appearing histologically normal. With the group that received only unheated oil, the aorta and perivascular tissue appeared histologically normal as shown in Fig 8 B. The photomicrograph shows cross sections and longitudinal sections of the aorta with surrounding fibrofatty tissue and a lymph node. The lumen of the aorta has a collection of red blood cells. The wall of the aorta appears histologically normal. Sections presented in Fig 8 C are representative cross sections from rats that received only heated oil. The section shows arteries adjacent to an airway with cartilage within its wall and lined by respiratory type epithelium. There is an adjacent lymph node in a fibrofatty stroma. The tissue is poorly dehydrated. There are chronic inflammatory changes within the wall of some small to medium sized venules close to the larger arteries. Though the aorta and the perivascular tissue are normal, there is isolated perivascular inflammation. Again, Fig 8 D show a cross section of arteries adjacent to an airway with cartilage within its wall and lined by respiratory type epithelium. There is an adjacent lymph node in a fibrofatty stroma. The tissue is poorly dehydrated with chronic inflammatory changes within the perivascular fat. This is normal aorta and perivascular tissue. Section of from the aortic tissue of rats treated with low dose celecoxib (CXB 5 mg/kg) as presented on Fig 8 E show cross sections of the aorta with surrounding fibrofatty tissue and a lymph node. The lumen of the aorta has a collection of red blood cells. The wall of the aorta appears histologically normal. The periaortic fat has cellular areas containing vacuolated adipocytes with features of blast cells. Similarly, a cross section of the aorta of rats treated with high dose celecoxib (CXB 10 mg/kg) showed the lumen of the aorta has a collection of red blood cells with surrounding fibrofatty tissue. The wall of the aorta appears histologically normal. The periaortic fat has cellular areas containing vacuolated adipocytes with features of blast cells. The aorta and the perivascular tissues appear histologically normal (Fig 8 F).

**Figure 8:**
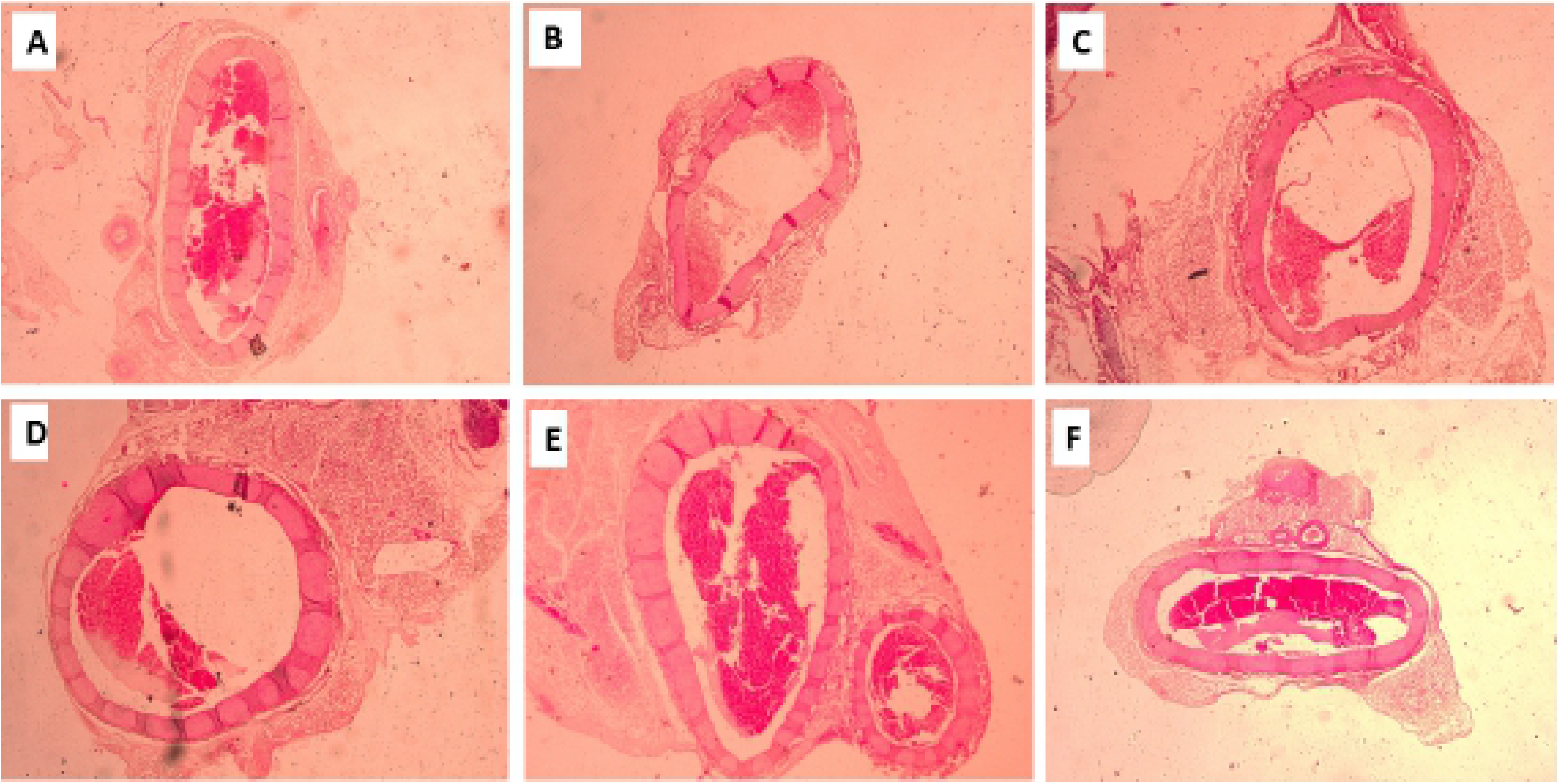
Photomicrograph of section of aorta of representative rat in (A) naïve control group (B) unheated oil treated group (C) heated oil only treated group as negative control, (D) heated oil in addition to atorvastatin 25 mg/kg (E) heated oil in addition to celecoxib 5 mg/kg and (F) heated oil in addition to celecoxib 10 mg/kg (H & E, ×40).

## Discussion

The global increase in the incidence of cardiovascular events continues to present a major public health issue because treatment remains suboptimal. Evidence abounds that lipid lowering therapy with statins (or ezetimibe in combination with a statin) contributes to reducing major adverse cardiovascular events. In spite of this, substantial risk of cardiovascular events remains even among patients receiving statin therapy whose LDL-C is <70 mg/dL [29, 30]. Alternative strategies are also required to lower lipids in patients who experience adverse effects on maximally tolerated statin therapy. These challenges call for innovations in the field of dyslipidaemia to address the several areas of unmet need [29].

Recent studies have demonstrated the association between increase in the expression of COX-2 and the development of metabolic disorders including obesity, diabetes mellitus, and non-alcoholic fatty liver disease (NAFLD). Studies have also shown that COX-2 activity not only has influence on insulin sensitivity [31], but also acts as pro-inflammatory mediator during the progression of NAFLD [32]. This latter has gained prominence as part of the possible mechanisms contributing to the protective effect of celecoxib against the development of NAFLD. Many studies suggested that celecoxib could attenuate liver steatosis and inflammation in NAFLD [33]. We also observed the ability of celecoxib to lower plasma cholesterol and attenuate hepatic lipid peroxidation in CCl_4_-mediated hepatotoxicity in rats [27]. In this study, we examined if the observed hypocholesterolaemic property of celecoxib in our previous study was unrelated to its hepatoprotective effect by evaluating its lipid lowering potential in rats fed with high fat (heated coconut oil) and without chemically-mediated induction of hepatic injury.

The results presented in Figure 1 describe the effect of celecoxib (CXB 5 and 10 mg/kg) and atorvastatin (ATO 25 mg/kg) on hyperlipidaemia induced by heated coconut oil. Naïve control group received only normal saline (10 mL/kg) throughout the experiment, the negative control group received heated oil and normal saline (10 mL/kg) whereas another group received unheated oil and normal saline (10 mL/kg). The relative liver weight (liver-to-body weight ratio) of the group that received only heated oil was significantly (P < 0.05) higher compared to the naïve control and unheated oil group. This was, however, significantly (P<0.05) decreased by treatment with celecoxib and atorvastatin. The relative weights of other organs such as heart, kidney, lungs and spleen were not significantly affected as shown in Figure 1. Generally, when oil is subjected to high temperature heating, free radicals are generated [34]. This may lead to several pathological changes in some organs as seen in the significantly increased weight of the liver. This remarkable increase in liver-to-body weight ratio has been attributed to the ability of the oil to increase liver microsomal lipid composition resulting in fatty liver [35].

Earlier reports suggests that there is strong correlations (both positive and negative) between the haematological parameters and the different lipid parameters [36]. Despite this fact, with the exception of platelet count, none of the haematological parameters assessed in the study was significantly affected (Figures 2 and 3). There was, however, a remarkable (P<0.05) increase in the platelet count (thrombocytosis) observed the group treated with heated oil only compared to the naïve control. This increase was also observed in the unheated oil treated group, though not significant. Among the causes of thrombocytosis is oxidative stress, inflammation, trauma, heart attack, cancer and burns [37]. We observed that celecoxib, but not atorvastatin, decreased the platelet count, although this effect was not statistically significant. This effect could be attributed to the anti-inflammatory effect of celecoxib in addition to its ability to ameliorate oxidative stress as reported earlier by Ekor *et al*. [27].

Results presented in Figure 4 show alanine aminotransferase (ALT) and alkaline phosphatase (ALP) activities of rats fed with heated coconut oil were significantly (P<0.05) higher than those of the naïve control. This is an indication of a possible hepatocellular damage [38]. The ALT enzyme is distributed in many tissues, but higher levels are present in the liver with elevated serum levels found in hepatocellular disorders than in intrahepatic or extra-hepatic cholestatic disorders 65. This result also confirms the possible involvement of liver disease and hypercholesterolaemia [27, 38]. Low and high doses of celecoxib significantly (all P<0.01) reversed this effect and this could be pointing to an ameliorative or protective effect of celecoxib against liver dysfunction associated with hyperlipidaemia.

Furthermore, we observed that sub-chronic administration of heated oil produced a significant decrease in blood urea levels in the rats. This decrease was significantly reversed by celecoxib. Although not very common, a decrease in urea levels could reflect severe liver disease [39]. Rather obvious and significant (P<0.05 for all) was the increase in total cholesterol levels, triglyceride levels, LDL, and VLDL levels in the heated oil treated group as shown in Figure 5. Treatment with atorvastatin and celecoxib significantly (p<0.05) ameliorated this effect. Hyperlipidaemia with a noticeable increase of low-density lipoprotein (LDL) cholesterol levels is common in patients with chronic cholestatic liver disease [39]. Therefore, the corresponding increase in ALT is not surprising though an increase in AST should have been expected. Some lipoproteins (notably those containing apoprotein B-100) are retained in the sub-endothelial space, by means of a charge-mediated interaction with extracellular matrix and proteoglycans [40]. This allows reactive oxygen species to modify the surface phospholipids and unesterified cholesterol of the small LDL particles. Because of LDL oxidation, isoprostanes are formed [41]. Vasoconstriction in the setting of high levels of oxidized LDL appear to be associated with a reduced release of the vasodilator nitric oxide from the damaged endothelial wall as well as increased platelet aggregation and thromboxane release [38, 42]. The state of hypercholesterolaemia leads invariably to an excess accumulation of oxidized LDL within the macrophages, thereby transforming them into “foam” cells. The rupture of these cells can lead to further damage of the vessel wall due to the release of oxygen free radicals, oxidized LDL, and intracellular enzymes [42].

Abnormal production of some cytokines such as tumour necrosis factor (TNF)-α, interleukin-1-beta (IL-1b), soluble IL-2 receptor (sIL-2R), IL-6, and chemokine IL-8 have been implicated in the pathogenesis of various inflammatory and autoimmune diseases [43]. When the sera of animals treated with heated oil were tested for serum cytokines (IL-6 and TNF-α), levels were not significantly (P>0.05 for both) affected as shown in Figure 6. Though many *in vivo* studies have demonstrated that TNF-α and IL-6 are important components of the pro-inflammatory response [43], this was not observed in our study.

Since the liver was the only organ whose relative weight was significantly affected by the various treatments, it was expedient to conduct a histopatholigical study on them. This falls in line the recommendation of the Society for Pathology and Toxicology (STP) that organ weights should be interpreted alongside hisptopathology [44]. Results obtained from the histology of the tissues show a distorted architecture of the liver intense congestion of central veins. Also, a large area of confluent necrosis with bridging, demarcated by only surrounding residual fibrocollagenous meshwork and surrounding central veins was observed. The periportal vessels and the area surrounding the central vein showed intense chronic inflammatory changes. This is a case of massive necrosis with ductular reaction and hepatitis. Treatment with celecoxib and atorvastatin, however, reversed to a very large extent this damage. This corroborates with the findings on the effect of the test drugs on liver enzymes such as ALT and ALP which are important markers of liver damage [45].

Though the aorta of the rats that received heated oil only was normal to a large extent, there were isolated spots of perivascular inflammation as well as fibrofatty stroma of the surrounding tissues. These were resolved to a large extent following treatment with celecoxib and atorvastatin.

## Conclusion

Celecoxib exhibited an attenuating effect on hyperlipidaemia and liver injury associated with sub-chronic ingestion of high fat (heated oil) in rats. Overall, findings from this study suggest that celecoxib, in addition to its established anti-inflammatory property, may be of therapeutic benefit in dyslipidaemia or related metabolic diseases and their attendant complications.

## Acknowledgements

The authors gratefully acknowledge Messrs. John K. Afortude, Wisdom Ahlidja and Joseph Acqua-Mills, of the Department of Pharmacology, School of Medical Sciences, University of Cape Coast, Cape Coast, Ghana, for their technical support.

## Funding

This study was funded by the University of Cape Coast through the Directorate of Research, Innovation and Consultancy (DRIC) research grant (Grant reference: RSG/GRP/CoHAS/2018/102).

## Declaration of competing interests

The authors declare that they have no competing interests.

## Authors’ contributions

ME: involved in conception and design of study, data analysis, revision of final draft of manuscript for important intellectual content, and submitted final manuscript. PEOA: involved in design of study, data analysis, revision of draft manuscript, and approved final manuscript for submission. EO: involved in design of study, data analysis, revision of draft manuscript, and approved final manuscript for submission. RPB: involved in design of study, data analysis, revision of draft manuscript, and approved final manuscript for submission. ITH: contributed to study design, data collection and analysis, drafted the manuscript and approved final manuscript for submission. MAA: contributed to data collection and analysis, and approved final manuscript for submission. GO: contributed to animal handling and data collection. ESY: involved in study design, revision of draft manuscript, and approved final manuscript for submission. PKA: contributed to data analysis, revised and approved the final submission of the manuscript.

## References

1. Joo IW, Ryu JH, Oh HJ. The influence of Sam-Chil-Geun (Panax notoginseng) on the serum lipid levels and inflammations of rats with hyperlipidemia induced by poloxamer-407. Yonsei Med J. 2010; 51(4):504–10.

2. Mc Namara K, Alzubaidi H, Jackson JK. Cardiovascular disease as a leading cause of death: how are pharmacists getting involved? Integr Pharm Res Pract. 2019;8:1–11.

3. Kraakman MJ, Dragoljevic D, Kammoun HL, Murphy AJ. Is the risk of cardiovascular disease altered with anti-inflammatory therapies? Insights from rheumatoid arthritis. Clin Transl Immunol. 2016. 5(5);e84; doi:10.1038/cti.2016.31

4. Stone NJ, Robinson J, Lichtenstein AH., Bairey Merz CN, Lloyd-Jones DM, Blum CB, et al. ACC/AHA guideline on the treatment of blood cholesterol to reduce atherosclerotic cardiovascular risk in adults: a report of the American College of Cardiology/American Heart Association Task Force on Practice Guidelines. J Am Coll Cardiol. 2013;63(25 Pt B):2889–934.

5. Perk J, De Backer G, Gohlke H, Graham I, Reiner Z, Verschuren M. European Association for Cardiovascular Prevention and Rehabilitation (EACPR), ESC Committee for Practice, et al. European guidelines on cardiovascular disease prevention in clinical practice (version 2012). The Fifth Joint Task Force of the European Society of Cardiology and Other Societies on Cardiovascular Disease Prevention in Clinical Practice (constituted by representatives of nine societies and by invited experts). Eur Heart J. 2012;13(13):635–701.

6. WHO 2008. WHO. The World Health report. WHO; Geneva: 2008. Cardiovascular disease. Available from: www.who.int/cardiovascular.diseases/en/ [2019 September 9th]

7. Mbikay M. Therapeutic potential of Moringa oleifera leaves in chronic hyperglycemia and dyslipidemia: a review. Front Pharmacol. 2012;3:24.

8. Nelson RH. Hyperlipidemia as a risk factor for cardiovascular disease. Prim Care. 2012;40(1):195–211.

9. Jou J, Choi SS, Diehl AM. Mechanisms of disease progression in nonalcoholic fatty liver disease. Semin Liver Dis. 2008;28:370–9.

10. Liao XH, Cao X, Liu J, Xie XH, Sun YH, Zhong BH. Prevalence and features of fatty liver detected by physical examination in Guangzhou. World J Gastroenterol. 2013;19(32):5334–9.

11. Al Mamun A, Hashimoto M, Katakura M, Tanabe Y, Tsuchikura S, Hossain S, Shido O. Effect of dietary n-3 fatty acids supplementation on fatty acid metabolism in atorvastatin-administered SHR.Cg-Leprcp/NDmcr rats, a metabolic syndrome model. Biomed Pharmacother. 2017;85:372–9.

12. Rang HP, Ritter JM, Flower RJ, Henderson G. Rang & Dale’s Pharmacology E-Book: with STUDENT CONSULT Online Access. 2014. Elsevier Health Sciences.

13. Kourounakis AP, Victoratos P, Peroulis N, Stefanou N, Yiangou M, Hadjipetrou L, Kourounakis PN. Experimental hyperlipidemia and the effects of NSAIDS. Exp Mol Pathol. 2002;73:135–8.

14. Livshits A, Seidman DS. Role of Non-Steroidal Anti-Inflammatory Drugs in Gynecology. Pharmaceuticals. 2010;3:2082–9.

15. Ahmed S, Gul S, Zia-Ul-Haq M, Riaz M. Hypolipidemic effects of nimesulide and celecoxib in experimentally induced hypercholesterolemia in rabbits. Turk J Med Sci. 2015;45:277–83

16. Foxworthy PS, Perry DN, Eacho PI. Induction of peroxisomal beta-oxidation by nonsteroidal anti-inflammatory drugs. Toxicol Appl Pharmacol. 1993;118(2):271–4.

17. Gansevoort PT, Heeg JE, Dikkeschei FD, de Zeeuw D, de Jong PE, Dullaart RFF. Symptomatic antiproteinuric treatment decreases serum lipoprotein (A) concentration in patients with glomerular proteinuria. Nephrol Dial Transplant. 1994;9(3):244–50.

18. Stoller DK, Grorud CB, Michalek V, Buchwald H. Reduction of Atherosclerosis with Nonsteroidal Anti-Inflammatory Drugs. J Surg Res. 1993;54(1):7–11.

19. Dhawan V, Ganguly NK, Majumdar S, Chakavarti RN. Effect of indomethacin on serum lipids, lipoproteins, prostaglandins and the extent and severity of atherosclerosis in rhesus monkeys. Can J Cardiol. 1992; 8(3):306–12.

20. Belton O, Fitzgerald DJ. Cyclooxygenase isoforms and atherosclerosis. Expert Rev Mol Med. 2003;5(9);1–18.

21. FitzGerald GA, Patrono C. The coxibs, selective inhibitors of cyclooxygenase-2. N Engl J Med. 2001; 345(6): 433–42.

22. Graham DJ, Campen D, Hui R, Spence M, Cheetham C, Levy G, Shoor S, et al. Risk of acute myocardial infarction and sudden cardiac death in patients treated with cyclo-oxygenase 2 selective and non-selective non-steroidal anti-inflammatory drugs: nested case-control study. Lancet. 2005;365(9458):475–81.

23. Hudson M, Richard H, Pilote LDifferences in outcomes of patients with congestive heart failure prescribed celecoxib, rofecoxib, or non-steroidal anti-inflammatory drugs: population based study. Bri Med J. 2005;330(7504):1370.

24. McGettigan P, Henry D. Cardiovascular risk and inhibition of cyclooxygenase: a systematic review of the observational studies of selective and nonselective inhibitors of cyclooxygenase 2. JAMA. 296(13), 1633–644.

25. Kristensen LE, Jakobsen AK, Askling J, Nilsson F, Jacobsson LT. Safety of etoricoxib, celecoxib, and nonselective nonsteroidal antiinflammatory drugs in ankylosing spondylitis and other spondyloarthritis patients: a Swedish national population‐based cohort study. Arthritis Care Res. 2015;67(8):1137–49.

26. Friend M, Vucenik I, Miller M. Platelet responsiveness to aspirin in patients with hyperlipidaemia. Br Med J. 2003;326:82–3.

27. Ekor M, Odewabi AO, Kale OE, Adesanoye OA, Bamidele TO. Celecoxib, a selective cyclooxygenase-2 inhibitor, lowers plasma cholesterol and attenuates hepatic lipid peroxidation during carbon-tetrachloride–associated hepatotoxicity in rats. Drug Chem Toxicol. 2013;36(1):1–8.

28. Garber JC, Barbee RW, Bielitzki JT, Clayton L, Donovan J, Hendriksen C, et al. Guide for the care and use of laboratory animals. The National Academic Press, Washington DC 8: 220.

29. Graham I, Shear C, Graeff PD, Boulton C, Catapano AL, Stough WG, et al. New strategies for the development of lipid-lowering therapies to reduce cardiovascular risk. Eur Heart J. 2018;4:119–27.

30. Robinson JG, Huijgen R, Ray K, Persons J, Kastelein JJ, Pencina MJ. Determining when to add nonstatin therapy: a quantitative approach. J Am Coll Cardiol. 2016;68:2412–21.

31. Chen X, Xu S, Wei S, Deng Y, Li Y, Yang F, et al. Comparative Proteomic Study of Fatty Acid-treated Myoblasts Reveals Role of Cox-2 in Palmitate-induced Insulin Resistance. Sci Rep.2016; 6:21454.

32. Yu J, Ip E, Dela Peña A, Hou JY, Sesha J, Pera N, et al. COX-2 induction in mice with experimental nutritional steatohepatitis: Role as pro-inflammatory mediator. Hepatology. 2006; 43:826–836.

33. Chen J, Liu D, Bai Q, Song J, Guan J, Gao J, et al. Celecoxib attenuates liver steatosis and inflammation in non-alcoholic steatohepatitis induced by high-fat diet in rats. Mol Med Rep. 2011;4:811–816.

34. Adam SK, Das S, Soelaiman IN, Umar NA, Jaarin K. Consumption of repeatedly heated soy oil increases the serum parameters related to atherosclerosis in ovariectomized rats. Tohoku J Exp Med. 2008;215(3):219–26.

35. de la Casa E, Perez-Gonzalez N, Sanchez-Bernal C, Llanillo M, Effects of dietary oil related to the toxic oil syndrome on the lipids of guinea pig liver microsomes. Lipids. 1995;30(6):575–9

36. Antwi-Baffour S, Kyeremeh R, Boateng SO, Annison L, Seidu MA. Haematological parameters and lipid profile abnormalities among patients with Type-2 diabetes mellitus in Ghana. Lipids Health Dis. 2018;17(1):1–9.

37. Torregrosa JM, Ferrer-Marin F, Lozano ML, Moreno MJ, Martinez C, Anton AI, et al. Impaired leucocyte activation is underlining the lower thrombotic risk of essential thrombocythaemia patients with CALR mutations as compared with those with the JAK2 mutation. Bri J Haematol. 2016;172(5):813–815.

38. Longo M, Crosignani A, Podda M. Hyperlipidemia in Chronic Cholestatic Liver Disease. Curr Treat Options Gastroenterol. 2001;4(2):111–4.

39. Waring WS, Stephen AFL, Robinson ODG, Dow MA, Pettie JM. Serum urea concentration and the risk of hepatotoxicity after paracetamol overdose. QJM. 2008;101(5):359–63.

40. Skalen K, Gustafsson M, Rydberg EK. Subendothelial retention of atherogenic lipoproteins in early atherosclerosis. Nature. 2002;417:750–4.

41. Reilly MP, Pratico D, Delanty N, DiMinno G, Tremoli E, Rader. Increased formation of distinct F2 isoprostanes in hypercholesterolemia. Circulation. 1998; 98(25):2822–8.

42. Ross R. The pathogenesis of atherosclerosis: a perspective for the 1990s. Nature. 1993;362(6423):801–9.

43. Evereklioglu C, Er H, Türköz Y, Cekmen M. Serum levels of TNF-alpha, sIL-2R, IL-6, and IL-8 are increased and associated with elevated lipid peroxidation in patients with Behçet’s disease. Mediators Inflamm. 2002;11(2):87–93.

44. Sellers RS, Mortan D, Michael B, Roome N, Johnson JK, Yano BL. Society of Toxicologic Pathology position paper: organ weight recommendations for toxicology studies. Toxicol Pathol. 2007;35(5):751–5.

45. Gaw A, Murphy M, Srivastava R, Cowan RA, O’Reilly DSJ. Clinical Biochemistry E-Book: An Illustrated Colour Text: Elsevier Health Sciences. 2013.

